# Survivin as a Multifaceted Oncogenic Driver and Therapeutic Target in Renal Cell Carcinoma

**DOI:** 10.1101/2025.09.02.673805

**Authors:** Shivani Tuli, Gabrielle Inserra, Yamato Murakami, Yongho Bae, Dave Gau

## Abstract

Renal cell carcinoma (RCC) is a heterogeneous malignancy in which clear cell RCC (ccRCC) represents the most aggressive subtype. Survivin (BIRC5), an inhibitor of apoptosis and key regulator of mitosis, is frequently overexpressed in RCC and associated with poor prognosis, yet its broader role in kidney cancer biology remains poorly defined. Here, we analyzed transcriptomic data from the TCGA-KIRC cohort and found that advanced-stage ccRCC exhibits widespread dysregulation of cell cycle pathways, with 1,484 genes upregulated and 479 genes downregulated in stage IV compared to stage I tumors. To define survivin’s functional contribution, we performed loss-of-function and pharmacologic inhibition studies in RENCA cells. Survivin knockdown or treatment with the small molecule inhibitor YM155 significantly reduced proliferation, S-phase entry, and Cyclin D1 expression, while also impairing both collective and single-cell migration. Beyond cell cycle control, survivin depletion induced notable changes in mitochondrial morphology and bioenergetics, including increased mitochondrial content coupled with reduced oxygen consumption, suggesting accumulation of dysfunctional mitochondria due to impaired clearance. Collectively, these findings identify survivin as a multifaceted oncogenic driver in RCC that integrates cell cycle progression, cytoskeletal organization, and mitochondrial homeostasis. By revealing survivin’s dual roles in proliferative and metabolic adaptation, this work highlights survivin as both a prognostic biomarker and a therapeutic vulnerability, supporting future strategies that combine survivin inhibition with metabolic or cell cycle-directed therapies for advanced kidney cancer.

## INTRODUCTION

Renal cell carcinoma (RCC) accounts for approximately 85% of all primary renal neoplasms and is the ninth most common cancer in the United States, with an estimated 80,980 new cases and 14,510 deaths projected in 2025 (1). Arising from renal tubular epithelial cells, RCC is a heterogeneous disease with several histologically and molecularly distinct subtypes. Clear cell RCC (ccRCC) is the most common subtype, accounting for over 75% of RCC cases, followed by papillary RCC (15-20%) and chromophobe RCC (5%). Among these subtypes, ccRCC has the worst clinical outcomes, with 30% of patients presenting with local recurrence or distant metastases after initial treatment. Consequently, the median survival for metastatic RCC is approximately 13 months, with a dismal 5-year survival rate of less than 10%. In contrast, survival rates are highly dependent on the stage at diagnosis, with a 78.1% overall 5-year survival rate (2).

Survivin’s dual role as an inhibitor of apoptosis and a regulator of mitosis is well-documented. It suppresses apoptosis by interacting with caspases-3, -7, and -9 and stabilizes mitochondrial integrity through its association with X-linked IAP (XIAP) and other apoptotic regulators (3, 4). Additionally, survivin functions as component of the chromosomal passenger complex (CPC), where it ensures proper chromosome alignment and segregation during mitosis by regulating microtubule dynamics (5, 6). However, emerging evidence suggests that survivin’s impact extends beyond these canonical roles to include regulation of early cell cycle progression, mitochondrial function, and metabolic reprogramming, processes that are critical for tumor progression but remain poorly understood in kidney cancer (7).

Mitochondria are central regulators of energy metabolism, apoptosis, and cellular homeostasis. In cancer cells, mitochondrial dynamics, specifically the processes of fission and fusion, are often dysregulated to support rapid proliferation and resistance to stress. Survivin has been implicated in modulating these mitochondrial function through localizing to the mitochondria and interacting with chaperones such as heat shock protein 90 (Hsp90) (8). These interactions stabilize mitochondrial integrity and prevent the release of pro-apoptotic factors like cytochrome c, thereby promoting cell survival under stress conditions (4, 9). Mechanistically, survivin inhibits mitophagy by blocking Parkin recruitment to dysfunctional mitochondria, leading to the accumulation of respiratory-defective mitochondria (10). Mitophagy inhibition forces cancer cells to rely on glycolysis for energy production, a hallmark of metabolic reprogramming known as the Warburg effect.

Despite growing recognition of survivin’s role in mitochondrial function and metabolism, the consequence of its knockdown in kidney cancer remain poorly understood. Furthermore, most studies have relied on standard cancer cell lines (e.g., HeLa or MCF-7), limiting their relevance to kidney cancer, a malignancy with unique metabolic dependencies. Addressing this gap is essential for advancing our understanding of survivin’s multifaceted roles in kidney cancer biology. In this study, we investigate survivin’s multifaceted role in renal carcinogenesis using a murine kidney cancer model system. Through survivin knockdown experiments, we uncover its critical involvement in mitochondrial dynamics and energy homeostasis, interconnected processes essential for cancer cell proliferation and migration. Survivin depletion induces systemic dysregulation across these pathways, impairing mitochondrial architecture, metabolic plasticity, and structural integrity. These disruptions culminate in suppressed proliferative capacity and compromised metastatic behavior.

Our work resolves a knowledge gap by establishing survivin as a central regulator of mitochondrial-structural coupling in kidney cancer. The coordinated collapse of bioenergetic and cytoskeletal networks upon survivin loss reveals an exploitable metabolic-structural vulnerability. This integrative perspective repositions survivin as a linchpin oncogenic hub, where its targeting could simultaneously disrupt tumor bioenergetics, growth signaling, and motility machinery. By bridging mitochondrial function with cellular architecture, these findings provide a framework for developing combination therapies that exploit survivin’s dual roles in renal carcinoma progression.

## Methods

### Cell Culture and siRNA Transfection

RENCA cells (ATCC, CRL-2947) were cultured in RPMI-1640 medium supplemented with 10% fetal bovine serum (FBS), 100 U/mL penicillin, and 100 μg/mL streptomycin. RENCA cells were transfected with either survivin-specific siRNA (target - CCGUCAGUGAAUUCUUGAA) or a non-targeting control siRNA using Lipofectamine RNAiMAX (Thermo Fisher, 13778075) according to the manufacturer’s instructions. The siRNAs were purchased from Thermo Fisher and used at a final concentration of 10 nM.

### YM-155 Treatment

After 48 hours of serum starvation in RPMI medium containing 1 mg/mL bovine serum albumin (BSA), RENCA cells were incubated in 10% serum-containing medium with 0.5 μM, 1 μM, or 2 μM YM155 (11490, Cayman) for 24 hours.

### Adenovirus Infection

Serum-starved RENCA cells were infected with either GFP (1060, Vector Biolabs) or BIRC5 (1611, Vector Biolabs) adenoviruses at a multiplicity of infection (MOI) of 25 or 50. Cells were incubated with the adenovirus for 24 hours prior to subsequent experiments.

### EdU Incorporation

RENCA cells were seeded onto 12-mm glass coverslips and incubated with 20 μM EdU (C10337, Thermo Fisher) for 9 or 24 hours. Following incubation, cells were fixed in 3.7% formaldehyde in PBS for 15 minutes. EdU incorporation assay was conducted using the Click-iT EdU Alexa Fluor 488 Imaging Kit (C10337, Thermo Fisher) according to the manufacturer’s instructions. Coverslips were mounted on microscope slides using DAPI-containing mounting medium (P36962, Thermo Fisher) and sealed with nail polish. Experiments were independently repeated four times, and five fields of view per coverslip were analyzed to determine the percentage of EdU-positive RENCA cells. EdU incorporation was quantified as fold change relative to control (DMSO or GFP) as previously described.

### Cell Proliferation and Scratch Assay

For cell proliferation assay, cells were plated per well in 24-well plates using triplicate determinations, with proliferation assessed after 72-96 hours by trypsonizing and counting number of cells. For scratch assay, RENCA cells were grown to a confluent monolayer in 24-well plates, and a straight scratch was made through the cell layer using a sterile 10 μL pipette tip. After scratching, the wells were gently washed with PBS to remove debris. Wound closure was monitored by capturing images of the same scratch area at 0 and 24 hours using an inverted microscope. The remaining wound area at 24 hours was quantified using ImageJ software, and migration was expressed as the percentage of wound closure relative to the initial scratch area.

### Single Cell Migration Assay

For single-cell migration experiments cells were plated in 24-well plates coated with type I collagen (Millipore) overnight. Where indicated, YM-155 (Cayman, Cat. No. 11490) was added to the culture medium (at 1 μM concentration) overnight prior to time-lapse imaging. Time-lapse images of randomly migrating cells were collected using a 10× objective for 120 min at 1-min time intervals using cellSens (Olympus Life Science) software. The centroid of the cell nucleus was tracked using ImageJ, and the average speed of migration was computed on a per-cell basis as before (11).

### Mitochondrial Morphology and Function Assessment

Mitochondrial network morphology was visualized by immunofluorescence staining of TOM20, a mitochondrial outer membrane protein. Cells were fixed with 4% paraformaldehyde for 5 minutes and permeabilized with 0.1% Triton X-100. Samples were incubated with a primary antibody against TOM20 (1:200 dilution, D8T4N, Cell Signaling Technology) followed by an Alexa Fluor 647-conjugated secondary antibody (1:100, 111-607-003, Jackson Immuno). Cells were counterstained with ActinRed555 ReadyProbes (R37112, Thermo Fisher) and DAPI. Fluorescence images were taken with either 60× objective on Olympus with Cicero confocal microscopes. All images were background subtracted using ImageJ software for quantitative analysis. Mitochondrial respiration analysis was performed using an Agilent Seahorse XF Analyzer. Briefly, RENCA cells were seeded into a XFmini 8-well cell culture microplate (Agilent) at 50,000 cells/well. Measurements were performed in a Seahorse XF Analyzer (Agilent). For the Seahorse Cell Mito Stress Test profile, cells were incubated for 1 hour at 37°C in XF Seahorse medium supplemented with glucose, glutamine, and sodium pyruvate and without CO_2_. Oxygen consumption rate (OCR) was measured under standard conditions and after the addition of 1 μmol/L oligomycin, 1.5 μmol/L FCCP, and 1 μmol/L rotenone/1 μmol/L Antimycin A.

### Gene Expression Analysis

RNA sequencing data and associated clinical annotations for kidney renal clear cell carcinoma (KIRC) were obtained from The Cancer Genome Atlas (TCGA). Only patient data annotated with stage I or stage IV were included in the analysis, patients with intermediate stages (II-III) were excluded. Raw count data from stage I and stage IV tumors were normalized and the DESeq2 package in R was used to identify differentially expressed genes (DEGs) between the tumor stages. DEGs were defined as those with a log_2_foldchange>1 and a Benjamini-Hochberg adjusted p-value (FDR) <0.05 (14).

### Clustering Analysis

Cytoscape’s String application was used to create a network of a protein-protein interactions (PPI) for the upregulated DEGs(16). Nodes (DEGs) with a degree of 2 or less were excluded to minimize small, disconnected clusters. The resulting network was clustered using kmeans clustering, with nine designated clusters, to create distinct regions of gene interactions. The clusters were named by their most enriched GO term, and the network was displayed on the String online server for visualization purposes.

### Protein Extraction and Immunoblotting

Total cell lysates were collected as follows. RENCA cells were washed twice with cold 1x DPBS and then incubated with 5X sample buffer (250 mM Tris-HCl [pH 6.8], 10% sodium dodecyl sulfate, 50% glycerol, 0.02% bromophenol blue, and 10 mM 2-mercaptoethanol) for 2 minutes. Cells were then scraped 10 times using a cell scraper. Protein samples were denatured at 100°C, and equal amounts were fractionated on 6-15% SDS-PAGE gels followed by transfer electrophoretically onto polyvinylidene difluoride membranes using Bio-Rad’s Trans-Blot Turbo Transfer System. Membranes were blocked in 6% milk in 1× Tris-buffered saline with 0.1% Tween 20 (TBST) prior to antibody incubation. Primary antibodies against survivin (NB500-201, Novus Biologicals; 1:500) and cyclin D1 (sc-20044, Santa Cruz Biotechnology; 1:200), were incubated at room temperature for 1 hour and overnight at 4°C. Following incubation, membranes were washed three times with 1x TBST and then probed with secondary antibodies (HRP-conjugated Affinipure Goat Anti-Rabbit IgG; Cat. No. SA00001-2, Proteintech or HRP-conjugated Affinipure Goat Anti-Mouse IgG; Cat. No. SA00001-1, Proteintech) for 1 hour at room temperature. Membranes were washed three times with 1x TBST prior to imaging.

Antibody signals were detected using Clarity Western ECL Substrate (Cat. No. 170-5061, Bio-Rad) or Clarity Max Western ECL Substrate (Cat. No. 1705062, Bio-Rad). All image analyses were performed using Bio-Rad Image Lab Software, and protein levels of survivin and cyclin D1, were normalized to GAPDH or α-tubulin as loading controls.

### Statistical Analysis

All experiments were conducted with biological triplicates (n = 3) for each condition, unless otherwise stated. Data are presented as the mean ± standard deviation (SD). Statistical comparisons between two groups (e.g., control vs. survivin-knockdown) were made using unpaired two-tailed Student’s t-tests. A p-value < 0.05 was considered statistically significant for all analyses. GraphPad Prism (v8.0) was used for data plotting and statistical calculations.

## RESULTS

### Survivin Overexpression Predicts Aggressive RCC and Poor Patient Survival

Survivin is aberrantly overexpressed in all three key histological subtypes of human RCC (with the most significant increase of expression in clear cell renal cell carcinoma) (**Fig. 1A**). Given that ccRCC is the most abundant form of kidney cancer, we focused on this subtype and found that high expression of survivin (segregated by median expression) showed worse patient prognosis (hazard ratio = 1.8; log-rank *P* = 0.00027; **Fig. 1B**), underscoring its clinical relevance as a biomarker for poor outcomes. High survivin levels in patients were strongly associated with advanced disease features such as higher stage tumor (**Fig. 1C**). This finding is comparable to previous study showing RCC patient cohorts with high survivin expression had significantly shorter overall survival than those with low expression (5-year cancer-specific survival ∼43% vs ∼87%, *p* < 0.001), and in addition, multivariate Cox regression confirmed survivin as an independent prognostic factor (after controlling for stage and grade, high tumoral survivin conferred a greater hazard of death with hazard ratio > 2, *p* < 0.01) (17). These data suggests that survivin expression correlates with RCC aggressiveness and poor patient prognosis, supporting its utility as a biomarker for aggressive disease.

**Figure 1.**
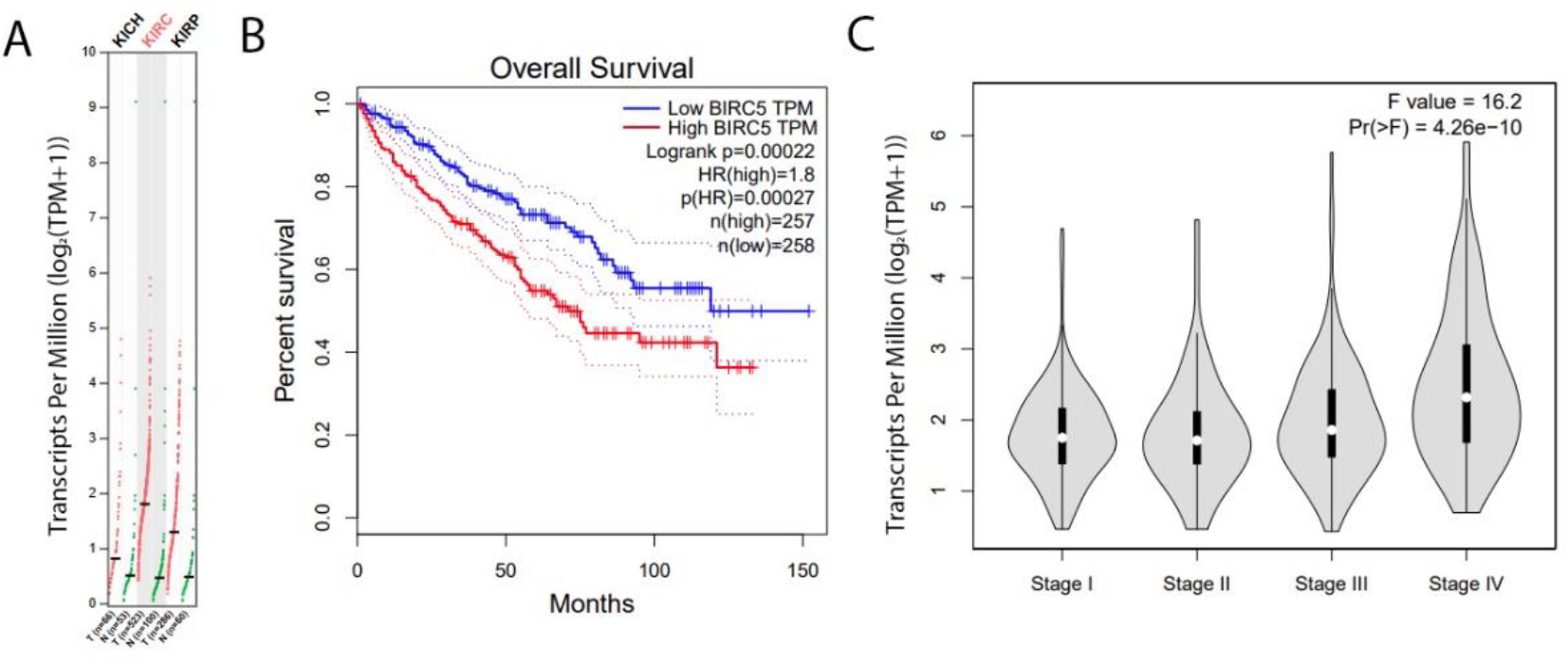
Survivin is transcriptionally upregulated in vast majority of human RCC. **A)** Relative survivin mRNA expression between tumors (T) of different histological subtypes of kidney cancer and normal kidney tissue (N). Patient sample numbers for different subtypes: chromophobe (T=66, N=53), clear cell (T=523, N=100), and papillary (T=286, N=60)). **B)** Kaplan-meier survival curve of TCGA KIRC dataset separated by median expression of survivin. **C)** Survivin expression plotted by different stages of ccRCC. Data were plotted on Gene Expression Profiling Interactive Analyses (GEPIA) platform (http://gepia.cancer-pku.cn/) that uses transcriptome data from the TCGA (the Cancer Genome Atlas) and the GTEx (Genotype Tissue Expression) databases.

### Transcriptomic Analysis of TCGA-KIRC Reveals Stage-Specific Dysregulation of Cell Cycle Pathways

To evaluate transcriptomic differences associated with disease progression, we analyzed publicly available TCGA clear cell renal cell carcinoma (TCGA-KIRC) data, focusing on stage I versus stage IV tumors. After quality control and filtering, differential expression analysis identified 479 downregulated and 1,484 upregulated genes. A PPI network was constructed to identify highly connected genes among the upregulated DEGs. Cytoscape’s String application was used to filter nodes by a degree ≤2, to exclude disconnected nodes and nodes with minimal interactions. Kmeans clustering identified 9 functionally distinct clusters among the upregulated. Clustering analysis revealed distinct patterns of enrichment among the upregulated DEGs, with the majority of significantly clustered genes mapping to pathways involved in cellular immune responses and cell cycle regulation (**Fig 2**). The cluster identified for its functional role in mitotic cell cycling forms a highly interconnected gene network of key cell cycle regulators, including survivin (BIRC5). Other functionally distinct clusters in the PPI network included genes associated with cell fate commitment, hormonal activity, ECM structure, gated channel activity, drug metabolism, intermediate filament organization, and tumor antigens. These findings suggest that advanced-stage ccRCC is characterized by marked dysregulation of cell proliferation–related processes, underscoring the central role of aberrant cell cycle control in disease progression.

**Figure 2.**
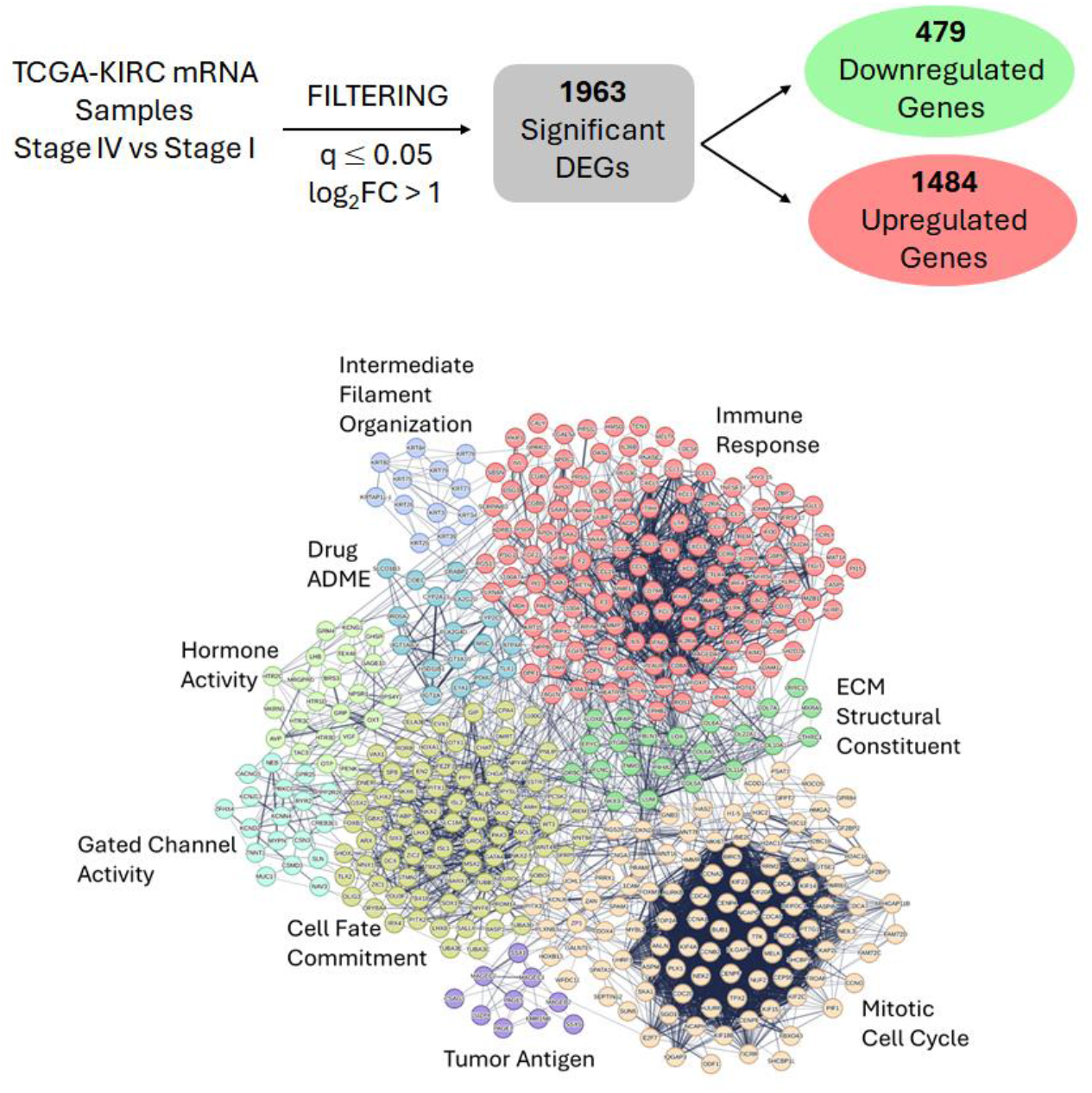
A highly interconnected cluster of cell cycle regulators is enriched in KIRC stage IV samples. Differential gene expression analysis was performed using TCGA-KIRC data comparing stage I and stage IV tumors. A total of 1,484 genes were upregulated and 479 genes were downregulated in stage IV tumors. Clustering analysis of these differentially expressed genes revealed a large, highly enriched cluster of genes associated with cell cycle regulation, highlighting enhanced proliferative signaling as a hallmark of advanced-stage ccRCC.

### Survivin inhibition decreases cell cycle progression and proliferation

Previous studies have shown that survivin is a key regulator of normal and cancer cell proliferation (4, 18-22). To directly assess survivin’s tumorigenic properties, we performed *in vitro* loss-of-function experiments in RENCA cells using siRNA. Strikingly, survivin knockdown (confirmed by western blot, **Fig. 3A**) caused a dramatic reduction in cancer cell proliferation compared to controls. Across replicate cell counting assays, survivin-deficient cells showed significantly lower growth rates (by ∼40% relative to control, *p*<0.0001, **Fig. 3B**). To further evaluate survivin’s role on RCC proliferation, we supplement our siRNA study with a small molecule inhibitor of survivin, YM155 (20, 23, 24). Cells were serum-starved to synchronize in G0, a quiescent state, and then incubated in growth medium containing YM155 or DMSO (vehicle control) for 24 hours. Immunoblotting of total cell lysates revealed that survivin inhibition significantly reduced the expression of Cyclin D1 (**Fig. 3C-E**), an early G1 cell cycle regulator. We also found that YM155 treatment significantly reduced S-phase entry, as assessed by EdU incorporation (**Fig. 3F**). To further determine if survivin overexpression affects S-phase entry, RENCA cells were infected with a wild-type survivin adenovirus or GFP control. Interestingly, survivin overexpression accelerated S-phase entry compared to GFP (**Fig. 3G**). Collectively, these findings suggest that survivin is required for cell cycle progression and proliferation in renal carcinoma.

**Figure 3.**
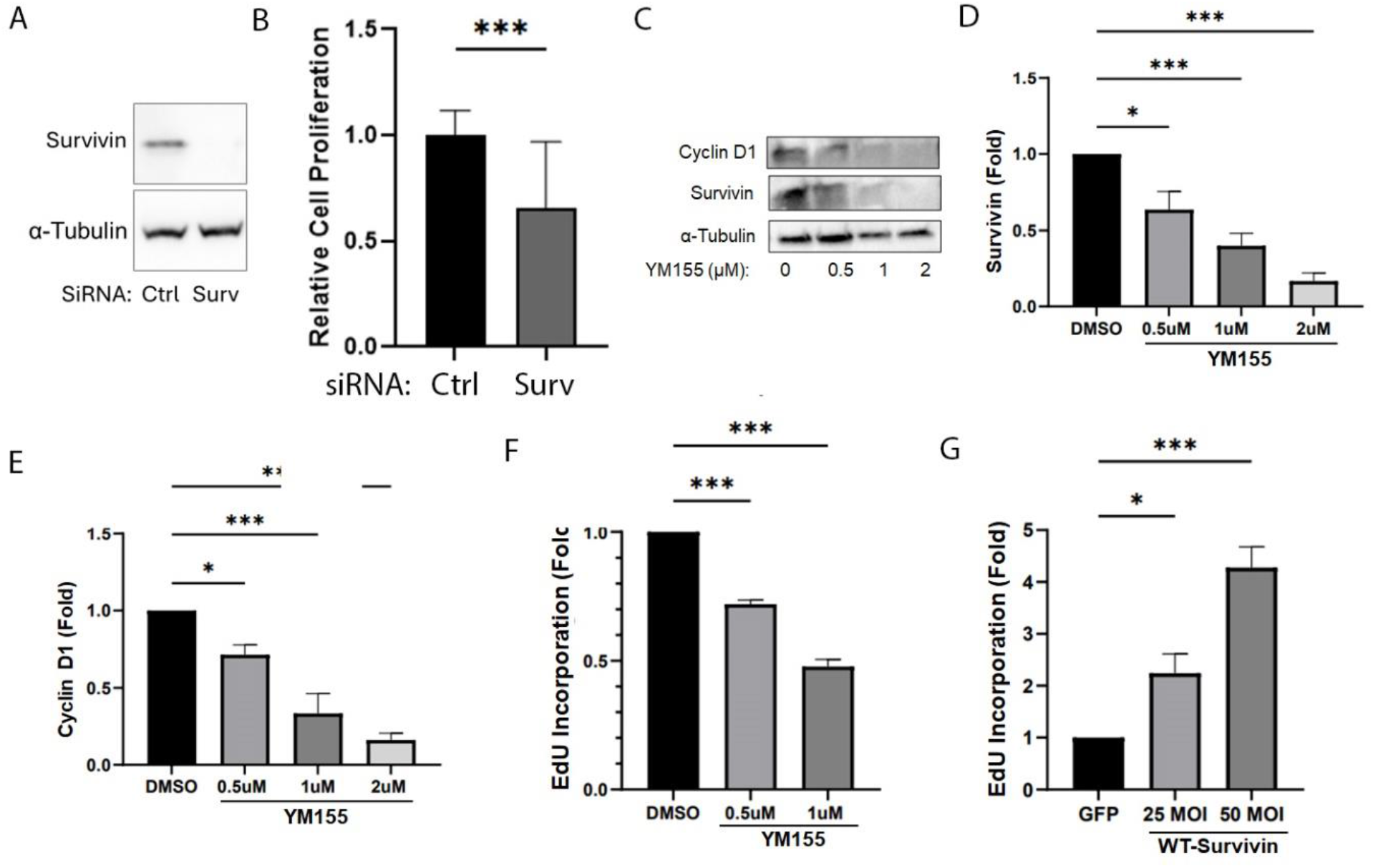
Survivin inhibition decreases proliferation and halts cell cycle progression. **A)** Immunoblot of survivin expression in control vs survivin siRNA treated RENCA cells. Tubulin serves as loading control. **B)** Bar graph depicting relative cell proliferation between ctrl vs survivin siRNA treated RENCA cells. Proliferation was assessed via cell counting. **C-E)** G0-synchronized RENCA cells were treated with 0.5μM, 1μM, and 2μM YM155 for 24 hours. Whole cell lysates were collected and analyzed with immunoblotting (C) to determine protein levels of Survivin (D) and Cyclin D1 (E). Expression levels were normalized to α-tubulin and displayed as a fold change to DMSO. n=4. G0-synchronized RENCA cells were treated with 1μM and 2μM YM155 for 24 hours. **F-G)** Expression levels were normalized to GAPDH and displayed as a fold change to DMSO. n=4. S-phase entry with YM155 treatment (F) and adenoviral overexpression (G) was assessed by EdU incorporation. Percentage of EdU positive cells was normalized to values from RENCA cells treated with DMSO (F) or GFP adenovirus (G). n=4. Data are means + SEMs. ^*^*p* < 0.05, ^**^*p* < 0.01, and ^***^*p* < 0.001

### Survivin inhibition decreases collective and single cell motility

Previous studies have shown that survivin is a key regulator of normal and cancer cell migration (25, 26). We first performed a scratch assay by plating control or survivin siRNA-treated RENCA cells to confluency. After 24 hours, survivin siRNA treated cells closed the scratch ∼50% less compared to control siRNA treated cells, **Fig. 4A-B**). Since scratch assays may be complicated by proliferation effects, especially when performed over longer periods of time, we also tracked cell migration via time-lapse video single cell migration assay. Our experimental data shows survivin siRNA reduced migration by almost 80% (**Fig. 4C**) and that 1µM YM155 treatment reduced migration by roughly 70% (**Fig. 4D**). This data suggests that reducing survivin not only stunts RENCA cell proliferation but also hinders cell motility, which together would limit tumor expansion, aligning with the known pro-tumorigenic role of survivin.

**Figure 4.**
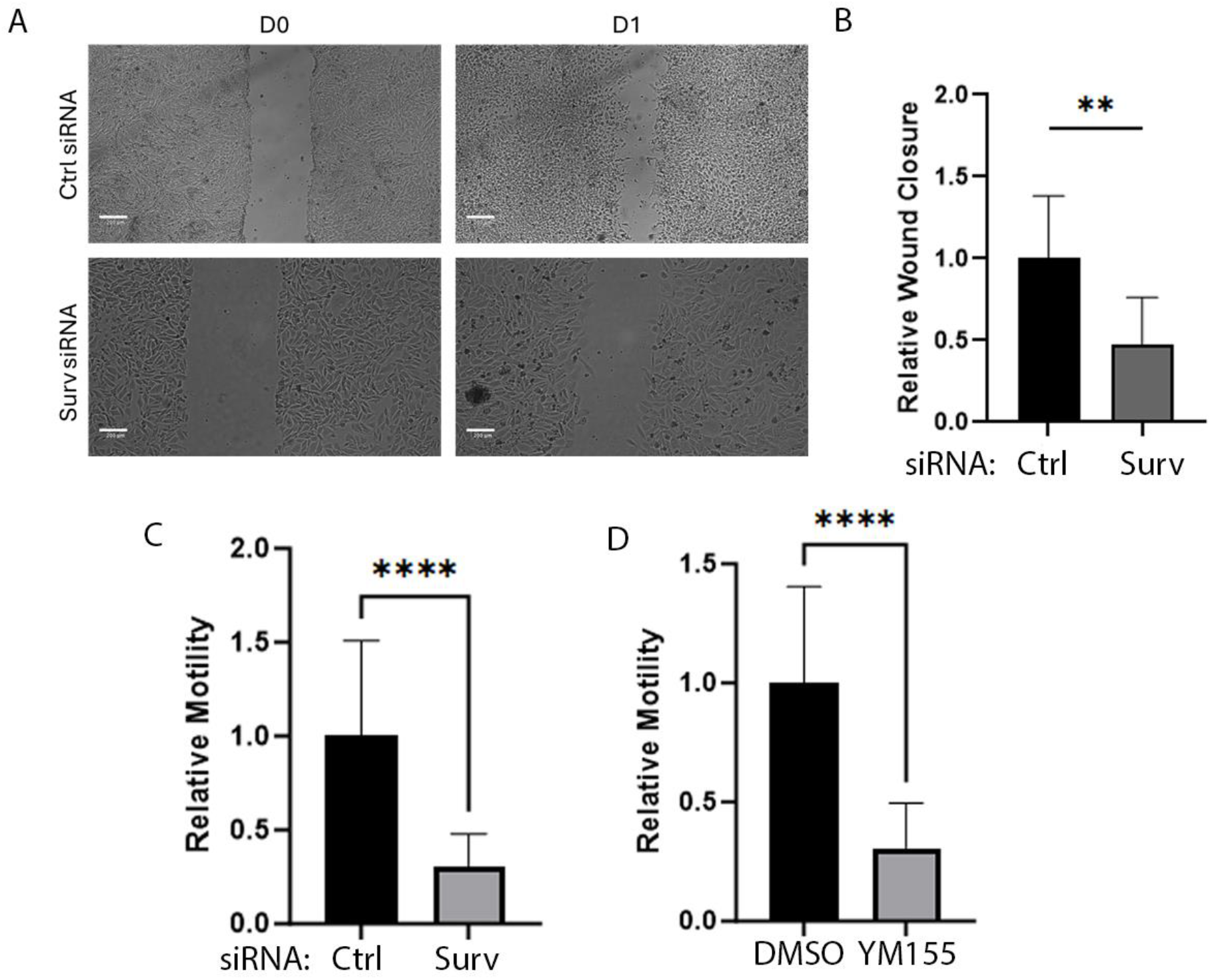
Loss of survivin decreased RENCA cell migration. **A)** Representative images of RENCA cells treated with ctrl or survivin siRNA in a scratch assay. Bar represents 200 µm. **B)** Bar graph depicting relative wound closure 24 hours after scratch. n = 3 experiments. **C-D)** Bar graph depicting relative speed of cells treated with ctrl vs. survivin siRNA or DMSO vs 1 µM YM155 via single cell migration assay. At least 60-80 cells were quantified from n = 3 experiments. Data are means + SD. **** p < 0.0001.

### Metabolic Dysfunction Induced by Loss of Survivin

In addition to functional deficits, immunostaining and image analysis demonstrated a reduction in mitochondrial content upon survivin knockdown. RENCA cells were immunostained to visualize mitochondria using a mitochondrial marker, TOM20, and images were captured on a widefield fluorescence microscope for quantitative analysis. Interestingly, Survivin knockdown cells showed increased fluorescent intensity of mitochondria staining and generally larger cell morphology (**Fig. 5A**). Quantification of the cell size and mitochondrial/cell area showed increased cell size (∼50% larger) and area of mitochondria relative (∼20% more mitochondria) to control siRNA treated cells (**Fig. 5B-C**). To examine the function of these mitochondria, we assessed oxygen consumption rate (OCR) and found a ∼30% decrease in basal OCR (**Supp. Fig. 1**). Together, these results indicate that loss of survivin expression further compromises the bioenergetic function of mitochondria in RENCA cells.

**Figure 5.**
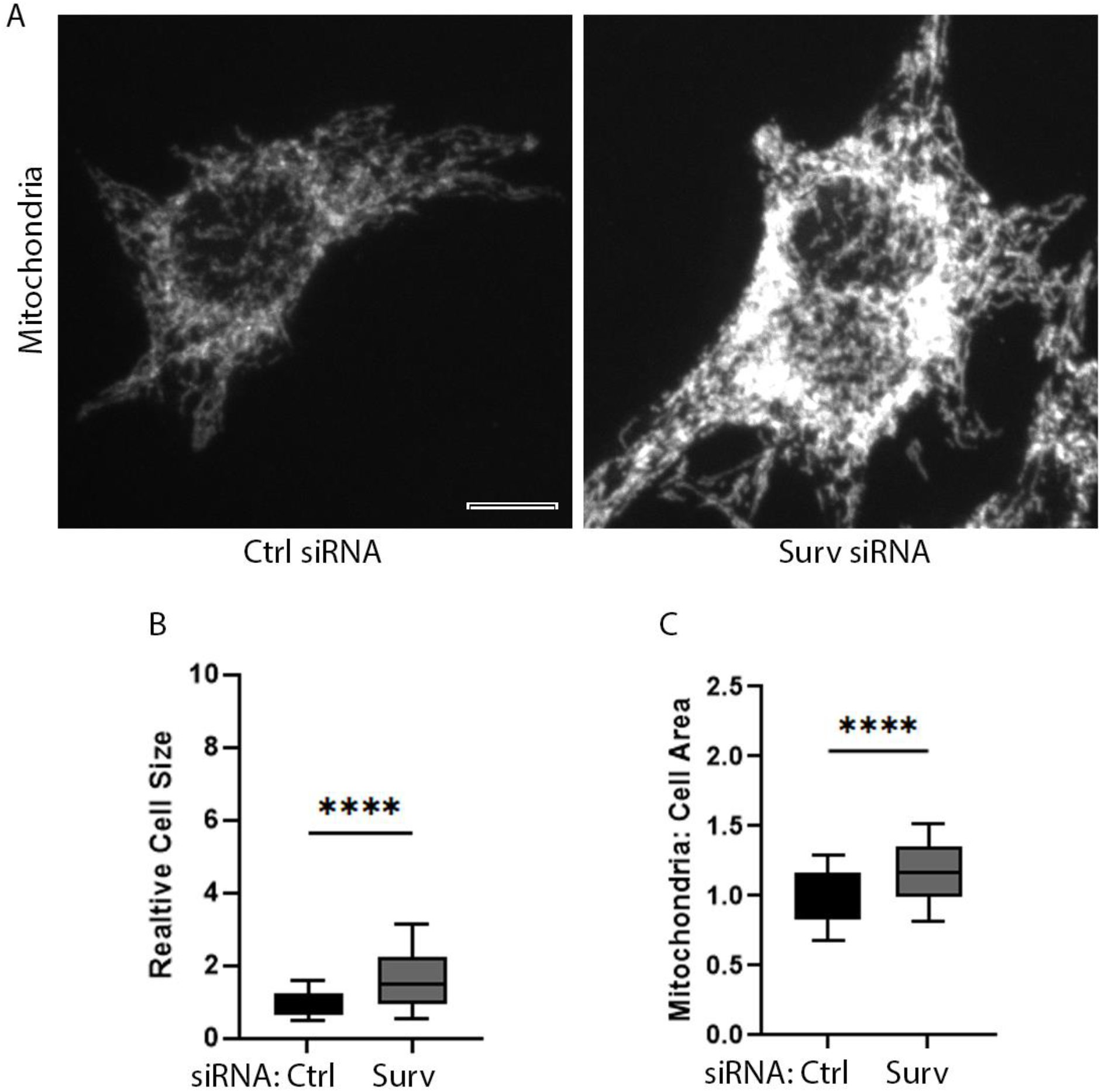
Survivin knockdown increases cell size and mitochondria content. **A)** Representative images of RENCA cells stained with Tom20 to visualize mitochondria. Bar represents 10 µm. **B-C)** Box and whisker plots depicting relative cell size and mitochondria/cell area between ctrl and survivin siRNA treated RENCA cells. At least 100-120 cells were quantified from n = 3 experiments. The bar represents the median with the box representing the 1^st^ and 3^rd^ quartile. Whiskers depict the 10^th^ and 90^th^ percentile.

## Discussion

Survivin is widely recognized for its canonical roles as an inhibitor of apoptosis and a regulated of mitotic progression, but our findings expand its functional repertoire in ccRCC. We demonstrate that survivin is not only indispensable for cell cycle progression and proliferation but also plays a critical role in maintaining mitochondria structure, bioenergetic efficiency, and cell migration. This suggests that survivin may serve as an important hub that coordinates the tumorigenesis of ccRCC. An interesting observation from our study was the increase of mitochondrial staining intensity and cell size after survivin knockdown despite a decrease in OCR. This suggests accumulation of dysfunctional mitochondria rather than enhanced biogenesis of mitochondria, potentially due to impaired clearance or mitophagy. This interpretation aligns with prior work implicating survivin in the inhibition of Parkin-mediated mitophagy (10). In RCC, this disruption may promote the persistence of structurally abundant yet functionally compromised mitochondria, exacerbating bioenergetic insufficiency and sensitizing tumor cells to metabolic stress. Follow-up studies assessing mitophagy and mitochondrial turnover will be important to confirm this mechanism.

Our data also emphasize survivin’s broader oncogenic functions in driving cell migration. The observation that both survivin knockdown and pharmacologic inhibition with YM155 substantially impaired RENCA cell motility suggests survivin links proliferative and cytoskeletal programs, consistent with other findings (32). Such dual regulation reinforces survivin’s role as a facilitator of tumor expansion and metastatic potential, consistent with its strong association with advanced tumor stage and poor prognosis in patient datasets. Therapeutically, these results underscore the promise of targeting survivin in RCC. While survivin inhibitors such as YM155 have shown preclinical efficacy, their clinical success has been limited, in part due to delivery challenges and activation of compensatory survival pathways (33). Our findings suggest that survivin inhibition could be more effective in rational combination regimens. For example, pairing survivin inhibitors with metabolic therapies (i.e. glycolysis or OXPHOS modulators) may exploit the mitochondrial vulnerabilities unmasked by survivin depletion. In addition, survivin’s role in sustaining proliferation and motility raises the possibility of synergistic benefit when combined with cell cycle inhibitors or immunotherapies. Survivin expression may also serve as a dual biomarker, identifying aggressive disease and predicting therapeutic response.

This study has limitations. Functional analyses were performed in RENCA cells, and validation across additional RCC models, including patient-derived samples, will be required to strengthen translational relevance. Our work was also limited to *in vitro* settings, and *in vivo* studies are essential to define the systemic impact of survivin inhibition on tumor progression and host metabolism. Finally, RCC is a heterogeneous disease, and survivin’s role may differ across subtypes and stages, necessitating broader investigation of context-specific functions. In summary, our findings position survivin as a multifaceted oncogenic driver in RCC that integrates cell cycle regulation, cytoskeletal organization, and mitochondrial homeostasis. Survivin depletion dismantles these interconnected networks, leading to cell cycle arrest, impaired migration, and bioenergetic collapse. By revealing survivin as a linchpin that bridges proliferative and metabolic programs, this work not only deepens our understanding of RCC biology but also highlights new opportunities for therapeutic intervention. Targeting survivin, particularly in combination with metabolic or immunomodulatory agents, represents a promising strategy to exploit the metabolic-structural vulnerabilities of RCC and improve outcomes for patients with this aggressive disease.

## Acknowledgement

The authors acknowledge funding from NIH grants R00CA267180 (Gau), R01HL163168 (Bae).

## Author Contributions

ST and GI performed experiments, conceived study design, analyzed data, and wrote the manuscript. YM performed experiments. YB and DG conceived study design, wrote the manuscript, and acquired funding.

## Data availability

All data are included in either main figures or supplementary information (SI)

## Figure Legends

**Supplemental Table 1**. Up-and down-regulated DEGs from TCGA analysis of KIRC dataset between stage I and stage IV cancers.

**Supplementary Figure 1.**
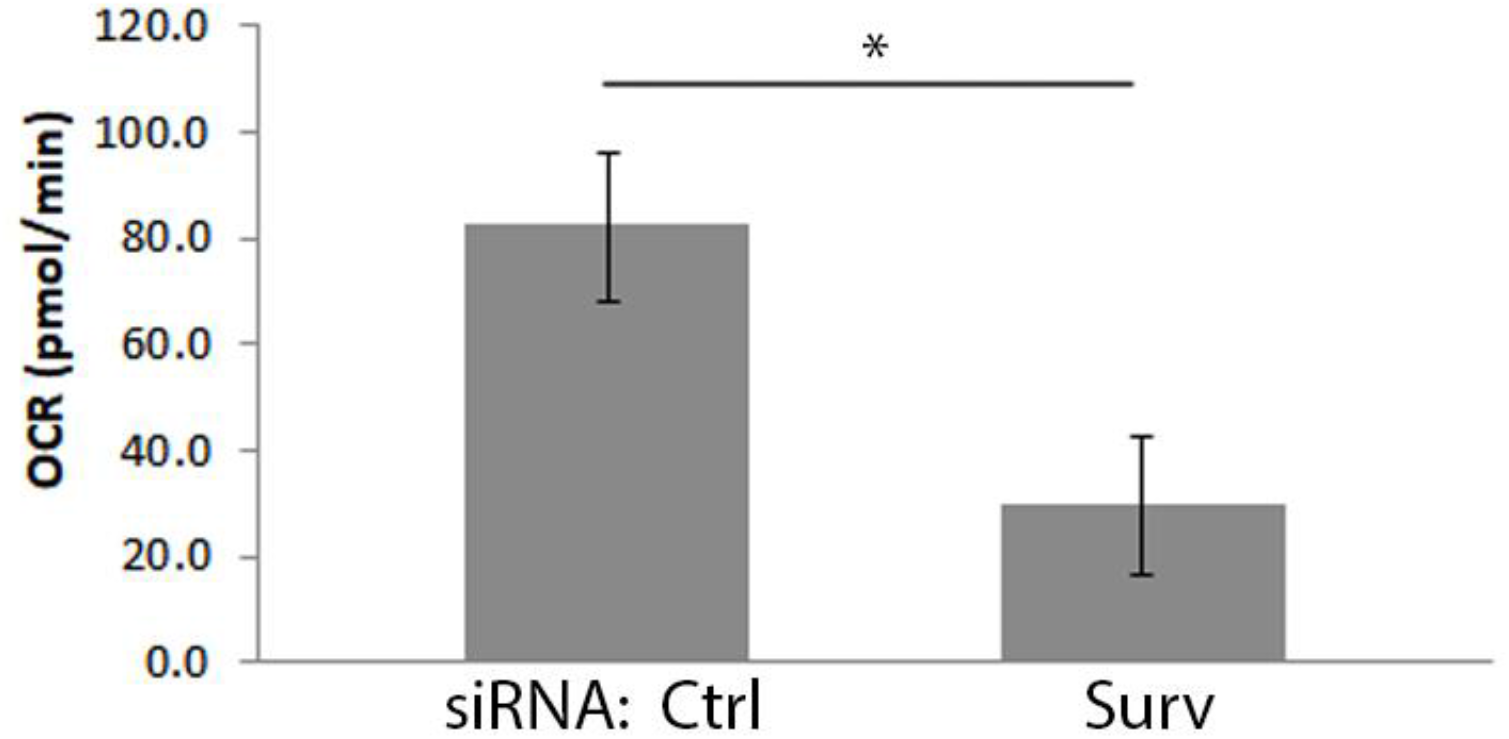
Loss of survivin reduces RENCA cell oxygen consumption rate. Bar graph showing OCR (pmol/min) of cells treated with survivin siRNA. n = 3 experiments. * p < 0.05.

## REFERENCES

1. Siegel, R. L., Kratzer, T. B., Giaquinto, A. N., Sung, H., andJemal, A. (2025) Cancer statistics, 2025 CA: a cancer journal for clinicians 75, 10–45 10.3322/caac.21871

2. Leibovich, B. C., Han, K. R., Bui, M. H., Pantuck, A. J., Dorey, F. J., Figlin, R. A. et al. (2003) Scoring algorithm to predict survival after nephrectomy and immunotherapy in patients with metastatic renal cell carcinoma: a stratification tool for prospective clinical trials Cancer 98, 2566–2575 10.1002/cncr.11851

3. Dohi, T., Beltrami, E., Wall, N. R., Plescia, J., andAltieri, D. C. (2004) Mitochondrial survivin inhibits apoptosis and promotes tumorigenesis J Clin Invest 114, 1117–1127 10.1172/JCI22222

4. Jaiswal, P. K., Goel, A., andMittal, R. D. (2015) Survivin: A molecular biomarker in cancer Indian J Med Res 141, 389–397 10.4103/0971-5916.159250

5. Knauer, S. K., Bier, C., Habtemichael, N., andStauber, R. H. (2006) The Survivin-Crm1 interaction is essential for chromosomal passenger complex localization and function EMBO reports 7, 1259–1265 10.1038/sj.embor.7400824

6. Vader, G., Kauw, J. J., Medema, R. H., andLens, S. M. (2006) Survivin mediates targeting of the chromosomal passenger complex to the centromere and midbody EMBO reports 7, 85–92 10.1038/sj.embor.7400562

7. Li, F., Aljahdali, I. A. M., Zhang, R., Nastiuk, K. L., Krolewski, J. J., andLing, X. (2021) Kidney cancer biomarkers and targets for therapeutics: survivin (BIRC5), XIAP, MCL-1, HIF1alpha, HIF2alpha, NRF2, MDM2, MDM4, p53, KRAS and AKT in renal cell carcinoma J Exp Clin Cancer Res 40, 254 10.1186/s13046-021-02026-1

8. Fortugno, P., Beltrami, E., Plescia, J., Fontana, J., Pradhan, D., Marchisio, P. C. et al. (2003) Regulation of survivin function by Hsp90 Proc Natl Acad Sci U S A 100, 13791–13796 10.1073/pnas.2434345100

9. Hennigs, J. K., Minner, S., Tennstedt, P., Loser, R., Huland, H., Klose, H. et al. (2020) Subcellular Compartmentalization of Survivin is Associated with Biological Aggressiveness and Prognosis in Prostate Cancer Sci Rep 10, 3250 10.1038/s41598-020-60064-9

10. Townley, A. R., andWheatley, S. P. (2020) Mitochondrial survivin reduces oxidative phosphorylation in cancer cells by inhibiting mitophagy J Cell Sci 133, 10.1242/jcs.247379

11. Gau, D., Veon, W., Shroff, S. G., andRoy, P. (2019) The VASP-profilin1 (Pfn1) interaction is critical for efficient cell migration and is regulated by cell-substrate adhesion in a PKA-dependent manner J Biol Chem 294, 6972–6985 10.1074/jbc.RA118.005255

12. Bolger, A. M., Lohse, M., andUsadel, B. (2014) Trimmomatic: a flexible trimmer for Illumina sequence data Bioinformatics 30, 2114–2120 10.1093/bioinformatics/btu170

13. Dobin, A., Davis, C. A., Schlesinger, F., Drenkow, J., Zaleski, C., Jha, S. et al. (2013) STAR: ultrafast universal RNA-seq aligner Bioinformatics 29, 15–21 10.1093/bioinformatics/bts635

14. Love, M. I., Huber, W., andAnders, S. (2014) Moderated estimation of fold change and dispersion for RNA-seq data with DESeq2 Genome Biol 15, 550 10.1186/s13059-014-0550-8

15. Reimand, J., Kull, M., Peterson, H., Hansen, J., andVilo, J. (2007) g:Profiler--a web-based toolset for functional profiling of gene lists from large-scale experiments Nucleic Acids Res 35, W193–200 10.1093/nar/gkm226

16. Szklarczyk, D., Franceschini, A., Wyder, S., Forslund, K., Heller, D., Huerta-Cepas, J. et al. (2015) STRING v10: protein-protein interaction networks, integrated over the tree of life Nucleic Acids Res 43, D447–452 10.1093/nar/gku1003

17. Parker, A. S., Kosari, F., Lohse, C. M., Houston Thompson, R., Kwon, E. D., Murphy, L. et al. (2006) High expression levels of survivin protein independently predict a poor outcome for patients who undergo surgery for clear cell renal cell carcinoma Cancer 107, 37–45 10.1002/cncr.21952

18. Mousso, T., Pham, K., Drewes, R., Babatunde, S., Jong, J., Krug, A. et al. (2025) Survivin in cardiovascular diseases and its therapeutic potential Vascul Pharmacol 159, 107475 10.1016/j.vph.2025.107475

19. Biber, J. C., Sullivan, A., Brazzo, J. A., 3rd, Heo, Y., Tumenbayar, B. I., Krajnik, A. et al. (2023) Survivin as a mediator of stiffness-induced cell cycle progression and proliferation of vascular smooth muscle cells APL Bioeng 7, 046108 10.1063/5.0150532

20. Guo, H., Wang, Y., Song, T., Xin, T., Zheng, Z., Zhong, P. et al. (2015) Silencing of survivin using YM155 inhibits invasion and suppresses proliferation in glioma cells Cell Biochem Biophys 71, 587–593 10.1007/s12013-014-0238-4

21. Chen, X., Duan, N., Zhang, C., andZhang, W. (2016) Survivin and Tumorigenesis: Molecular Mechanisms and Therapeutic Strategies J Cancer 7, 314–323 10.7150/jca.13332

22. Mobahat, M., Narendran, A., andRiabowol, K. (2014) Survivin as a preferential target for cancer therapy Int J Mol Sci 15, 2494–2516 10.3390/ijms15022494

23. Cheng, X. J., Lin, J. C., Ding, Y. F., Zhu, L., Ye, J., andTu, S. P. (2016) Survivin inhibitor YM155 suppresses gastric cancer xenograft growth in mice without affecting normal tissues Oncotarget 7, 7096–7109 10.18632/oncotarget.6898

24. Sasaki, R., Ito, S., Asahi, M., andIshida, Y. (2015) YM155 suppresses cell proliferation and induces cell death in human adult T-cell leukemia/lymphoma cells Leuk Res 39, 1473–1479 10.1016/j.leukres.2015.10.012

25. Nabzdyk, C. S., Lancero, H., Nguyen, K. P., Salek, S., andConte, M. S. (2011) RNA interference-mediated survivin gene knockdown induces growth arrest and reduced migration of vascular smooth muscle cells Am J Physiol Heart Circ Physiol 301, H1841–1849 10.1152/ajpheart.00089.2011

26. Mehrotra, S., Languino, L. R., Raskett, C. M., Mercurio, A. M., Dohi, T., andAltieri, D. C. (2010) IAP regulation of metastasis Cancer Cell 17, 53–64 10.1016/j.ccr.2009.11.021

27. Helmke, C., Becker, S., andStrebhardt, K. (2016) The role of Plk3 in oncogenesis Oncogene 35, 135–147 10.1038/onc.2015.105

28. Wang, B., Lan, T., Xiao, H., Chen, Z. H., Wei, C., Chen, L. F. et al. (2021) The expression profiles and prognostic values of HSP70s in hepatocellular carcinoma Cancer Cell Int 21, 286 10.1186/s12935-021-01987-9

29. Zhang, H., Duan, J., Qu, Y., Deng, T., Liu, R., Zhang, L. et al. (2016) Onco-miR-24 regulates cell growth and apoptosis by targeting BCL2L11 in gastric cancer Protein Cell 7, 141–151 10.1007/s13238-015-0234-5

30. Wu, H., Li, Z. X., Fang, K., Zhao, Z. Y., Sun, M. C., Feng, A. Q. et al. (2024) IGF-1-mediated FOXC1 overexpression induces stem-like properties through upregulating CBX7 and IGF-1R in esophageal squamous cell carcinoma Cell Death Discov 10, 102 10.1038/s41420-024-01864-0

31. Lee, J. T., Brafford, P., andHerlyn, M. (2008) Unraveling the mysteries of IGF-1 signaling in melanoma J Invest Dermatol 128, 1358–1360 10.1038/jid.2008.124

32. Guo, K., Huang, P., Xu, N. J., Xu, P., Kaku, H., Zheng, S. B. et al. (2015) A combination of YM-155, a small molecule survivin inhibitor, and IL-2 potently suppresses renal cell carcinoma in murine model Oncotarget 6, 21137–21147 DOI 10.18632/oncotarget.4121

33. Albadari, N., andLi, W. (2023) Survivin Small Molecules Inhibitors: Recent Advances and Challenges Molecules 28, 10.3390/molecules28031376

34. Xu, L., Hu, H., Zheng, L. S., Wang, M. Y., Mei, Y., Peng, L. X. et al. (2020) ETV4 is a theranostic target in clear cell renal cell carcinoma that promotes metastasis by activating the pro-metastatic gene FOSL1 in a PI3K-AKT dependent manner Cancer Lett 482, 74–89 10.1016/j.canlet.2020.04.002

35. Wei, Y., Han, S., Wen, J., Liao, J., Liang, J., Yu, J. et al. (2023) E26 transformation-specific transcription variant 5 in development and cancer: modification, regulation and function J Biomed Sci 30, 17 10.1186/s12929-023-00909-3

36. Sionov, R. V., Vlahopoulos, S. A., andGranot, Z. (2015) Regulation of Bim in Health and Disease Oncotarget 6, 23058–23134 10.18632/oncotarget.5492

37. Hagenbuchner, J., Kuznetsov, A. V., Obexer, P., andAusserlechner, M. J. (2013) BIRC5/Survivin enhances aerobic glycolysis and drug resistance by altered regulation of the mitochondrial fusion/fission machinery Oncogene 32, 4748–4757 10.1038/onc.2012.500

38. Yin, P., Xu, Q., andDuan, C. (2004) Paradoxical actions of endogenous and exogenous insulin-like growth factor-binding protein-5 revealed by RNA interference analysis J Biol Chem 279, 32660–32666 10.1074/jbc.M401378200

39. Sureshbabu, A., Okajima, H., Yamanaka, D., Tonner, E., Shastri, S., Maycock, J. et al. (2012) IGFBP5 induces cell adhesion, increases cell survival and inhibits cell migration in MCF-7 human breast cancer cells J Cell Sci 125, 1693–1705 10.1242/jcs.092882

40. Pantazi, E., Gemenetzidis, E., Trigiante, G., Warnes, G., Shan, L., Mao, X. et al. (2014) GLI2 induces genomic instability in human keratinocytes by inhibiting apoptosis Cell Death Dis 5, e1028 10.1038/cddis.2013.535

41. Carew, J. S., Espitia, C. M., Zhao, W., Mita, M. M., Mita, A. C., andNawrocki, S. T. (2015) Targeting Survivin Inhibits Renal Cell Carcinoma Progression and Enhances the Activity of Temsirolimus Mol Cancer Ther 14, 1404–1413 10.1158/1535-7163.MCT-14-1036

